# SARS-CoV-2 monoclonal antibody treatment followed by vaccination shifts human memory B cell epitope recognition suggesting antibody feedback

**DOI:** 10.1101/2023.11.21.567575

**Authors:** Camila H. Coelho, Nathaniel Bloom, Sydney I. Ramirez, Urvi M. Parikh, Amy Heaps, Scott F. Sieg, Alex Greninger, Justin Ritz, Carlee Moser, Joseph J. Eron, Judith S. Currier, Paul Klekotka, David A. Wohl, Eric S. Daar, Jonathan Li, Michael D. Hughes, Kara W. Chew, Davey M. Smith, Shane Crotty, the Accelerating COVID-19 Therapeutic Interventions and Vaccines–2/A5401 (ACTIV-2/A5401) Study Team

## Abstract

Therapeutic anti-SARS-CoV-2 monoclonal antibodies (mAbs) have been extensively studied in humans, but the impact on immune memory of mAb treatment during an ongoing immune response has remained unclear. Here, we evaluated the effect of infusion of the anti-SARS-CoV-2 spike receptor binding domain (RBD) mAb bamlanivimab on memory B cells (MBCs) in SARS-CoV-2–infected individuals. Bamlanivimab treatment skewed the repertoire of memory B cells targeting Spike towards non-RBD epitopes. Furthermore, the relative affinity of RBD memory B cells was weaker in mAb-treated individuals compared to placebo-treated individuals over time. Subsequently, after mRNA COVID-19 vaccination, memory B cell differences persisted and mapped to a specific defect in recognition of the class II RBD site, the same RBD epitope recognized by bamlanivimab. These findings indicate a substantial role of antibody feedback in regulating human memory B cell responses, both to infection and vaccination. These data indicate that mAb administration can promote alterations in the epitopes recognized by the B cell repertoire, and the single administration of mAb can continue to determine the fate of B cells in response to additional antigen exposures months later.

**SIGNIFICANCE STATEMENT:** Evaluating the therapeutic use of monoclonal antibodies during SARS-CoV-2 infection requires a comprehensive understanding of their impact on B cell responses at the cellular level and how these responses are shaped after vaccination. We report for the first time the effect of bamlanivimab on SARS-CoV-2 specific human memory B cells of COVID-19 infected humans receiving, or not, mRNA immunization.

## MAIN TEXT

During the COVID-19 pandemic, monoclonal antibodies (mAbs) were rapidly developed and tested for clinical safety and efficacy as prophylaxis and treatment for COVID-19, resulting in the emergency use authorization (EUA) from the U.S. Food and Drug Administration for several products and subsequent widespread clinical use in adults and children at risk for severe COVID-19 (1–3). ACTIV-2/A5401 was a multicenter phase 2/3 randomized controlled trial designed to evaluate the safety and efficacy of therapeutics for acute COVID-19 in non-hospitalized adults, including bamlanivimab (4). Bamlanivimab is a neutralizing human immunoglobulin (IgG)-1 that binds to the receptor binding domain (RBD) of the SARS-CoV-2 ancestral spike (S) protein, that was developed early in the pandemic. Similar to contemporaneously mAbs, bamlanivimab was shown to have antiviral activity in non-hospitalized adults with mild to moderate acute COVID-19 (2, 4, 5).

The effects of mAb treatment on immune responses remain incompletely characterized (6, 7). We recently showed that acute and memory CD4 and CD8 T cell responses were not measurably impacted by mAb treatment (8).

It has been observed that circulating antibodies can affect B cell responses (9–13), in a process known as antibody feedback. There is heightened interest in understanding the biology of antibody feedback, as recent data indicates it may be a more influential immunological process than previously recognized, and it may be relevant in important biomedical contexts ranging from mAb therapeutics (7) to imprinting challenges of universal flu or universal sarbecovirus vaccines (14–18). High affinity antibodies with singular specificity can block the recruitment of naïve B (10) and IgM^+^ B cells (9, 11). Antibody-mediated suppression is dose-dependent, and high concentrations of IgG can suppress a primary antibody response (11, 19). This effect can be mediated by any isotype of mAbs (9). There is also evidence that antibody feedback can influence germinal centers (13, 19).

Roles of antibody feedback in humans have been more challenging to define. A recent report studied a cocktail containing two SARS-CoV-2 RBD-targeting mAbs given to uninfected healthy participants as a safety study (7). Some participants subsequently received two doses of COVID-19 mRNA vaccines. The affinities of antibodies (Abs) from B cells isolated from the mAb treated group were significantly lower than those receiving vacination only. The response was also characterized by a reduced frequency of epitope-specific antibodies, demonstrating an antibody feedback or epitope masking phenomenon, shifting the B cell response away from the RBD sites bound by the therapeutic mAbs (7). Notably, mAbs obtained from people receiving mAb treatment were almost entirely unable to neutralize 7 different strains of SARS-CoV-2 pseudoviruses (7). It remains unclear if those outcomes are unique to mRNA vaccination, or whether similar outcomes would occur upon an infection. Furthermore, the impact of mAb treatment on B cell responses in humans during an ongoing infection has remained unclear, which has both clinical treatment and fundamental immunology implications.

In this study, we evaluated the effect of mAb administration during acute SARS-CoV-2 infection on memory B cell responses, and again later after mRNA vaccination, in participants from the ACTIV-2/A5401 trial of therapeutics for non-hospitalized adults with COVID-19.

## RESULTS

### c memory B cell development

Bamlanivimab 700 mg or placebo (saline) infusion was administered to participants with acute COVID-19 in the randomized ACTIV-2/A5401 trial (4, 20). A subset of participants was included in this analysis, selected based on the availability of longitudinal samples, as previously described (8). Clinical and demographic characteristics of the analysis cohort are described in **Table S1** and have been reported previously (8). In the analysis cohort (**Fig. 1A**), at 28 days post-treatment, IgG titers against SARS-CoV-2 Spike S1 and RBD were significantly increased in mAb recipients compared to the placebo group, as expected (**Fig. 1B-C**) (8). We analyzed the effect of bamlanivimab treatment on the frequency of SARS-CoV-2-specific memory B cells at 28 days post-treatment (**Fig. 1D-E, Fig. S1A-D**). No differences were detected in the frequency of MBCs binding to Spike or RBD (Spike MBCs and RBD MBCs, **Fig. 1F-G**). However, the proportion of Spike-specific B cells that bound RBD was significantly decreased in the treatment group compared to the placebo (**Fig. 1H**). Conversely, the frequency of non-RBD MBCs among Spike-specific MBCs (i.e., MBCs binding to other Spike domains such as the N-terminal domain or S2) was higher in the bamlanivimab group (**Fig. 1I**). The relative affinity of RBD MBCs, measured by the MFI ratio of probe to CD79b (21), trended towards being slightly lower at week 28 in the mAb treatment group (*p*=0.085, **Fig. 1J**). For Spike MBCs, the mAb treatment did not measurably alter BCR affinities overall (probe MFI/CD79b MFI. **Fig. S1A**). The frequencies of IgG^+^ and IgM^+^ RBD MBCs were comparable between groups (**Fig. 1K-L**). The development of IgA^+^ RBD MBCs was lower in bamlanivimab-treated individuals (**Fig. 1M**). We investigated whether mAb treatment altered MBC phenotypic development by analyzing selected surface markers (CXCR3, CXCR5, CD95, CD71, CD11c) together with CD27 and CD21 to quantify classical, atypical, and activated memory B cells. mAb treatment did not significantly alter the frequency of any phenotypic markers analyzed (**Fig. S3**).

**Figure 1.**
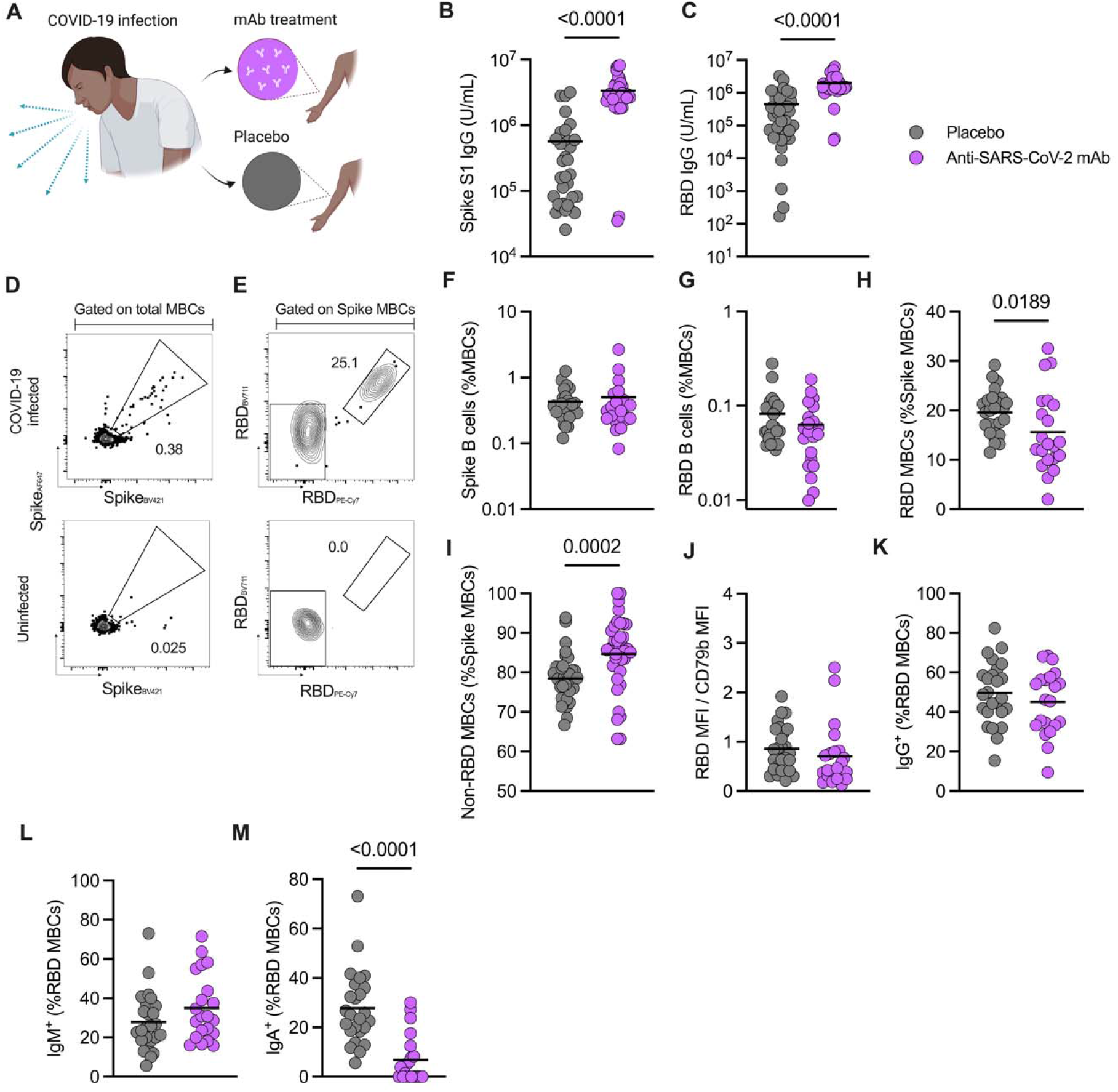
Effect of bamlanivimab after 28 days of treatment in COVID-19 cases. **(A)** SARS-CoV-2 infected people were treated with 700 mg intravenous dose of bamlanivimab **(B-C)** Spike S1 (B) and RBD (C) IgG titers in the placebo and mAb-treated groups **(D)** Representative gating strategy for spike-binding memory B cells (“MBCs”) in SARS-CoV-2 infected (top image) and uninfected humans (bottom image) (Refer also to **Fig. S5**). **(E)** Representative gating strategy for RBD-binding MBCs in SARS-CoV-2 infected (top image) and uninfected humans (bottom image). **(F-G)** Frequency of Spike (F) and RBD (G) MBCs among total memory B cells (IgD, CD27) **(H)** Frequency of RBD MBCs within the Spike MBCs population **(I)** Frequency of non-RBD MBCs within the Spike MBCs population **(J)** Affinity of RBD MBCs determined by the ratio MFI from the RBD probe (PE-CY7)/ MFI from immunoglobulin-associated beta (CD79b) **(K-M)** Frequency of RBD MBCs expressing IgG (K), IgM (L) or IgA (M). Means are shown. An unpaired, non-parametric Mann-Whitney test was used to compare the different groups. p-values are shown in the graphs and considered significant if lower than 0.05. **Sample size: treatment group n= 25 and placebo group n=21**

We then evaluated long-term effects of bamlanivimab on MBCs at 24 weeks (**Fig. 2, Fig. S1E-H**). While the frequency of Spike MBCs was not significantly different between bamlanivimab and placebo groups (**Fig. 2A**), the frequency of RBD MBCs was lower in the treatment group, of borderline statistical significance (*p* = 0.056. **Fig. 2B**). The percentage of Spike-specific MBCs that were RBD-specific differed to a greater extent between treatment and placebo groups at 24 weeks (6.5% *vs* 17.3%, *p*<0.0001, **Fig. 2C, 2D**) than at 4 weeks (15.6% *vs* 19.59%, *p*=0.0189, **Fig. 1H**). Furthermore, the relative affinity of RBD MBCs was significantly lower between treatment and placebo group after 24 weeks (*p*<0.0001, **Fig. 2E**). There was no difference between frequency of RBD MBCs expressing IgG in their surface (**Fig. 2F**) while the frequency of IgM+ MBCs was higher in the bamlanivimab group (**Fig. 2G**). The difference in IgA^+^ RBD MBCs frequency was less evident (*p*=0.054, **Fig. 2H**). These data demonstrate that mAb treatment significantly changed epitope recognition by B cells targeting RBD during a SARS-CoV-2 infection, with the defects becoming more apparent over time.

**Figure 2.**
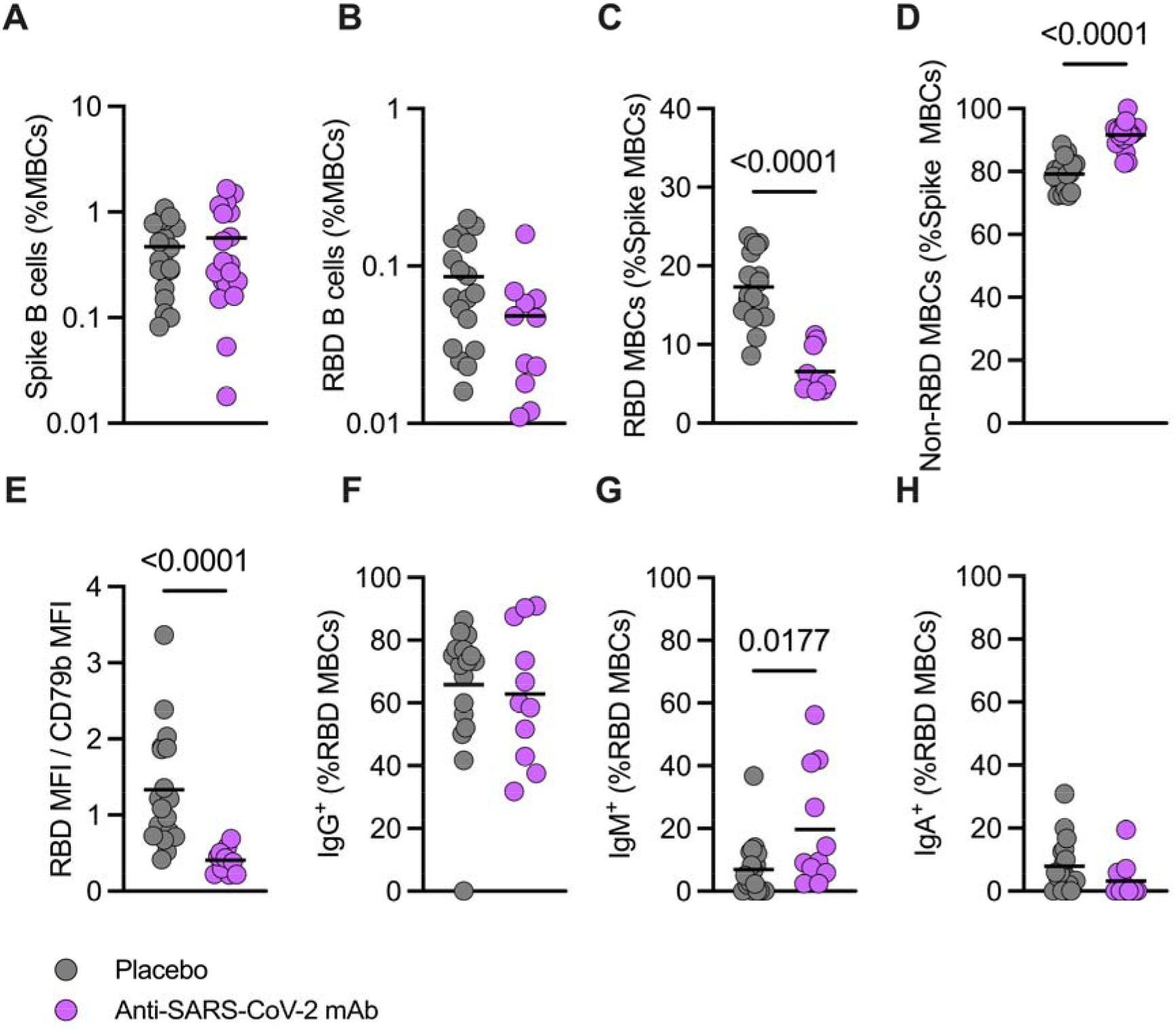
Effect of bamlanivimab after 24 weeks of treatment in COVID-19 infected adults. **(A-B)** Frequency of Spike (A) and RBD (B) MBCs among total memory B cells (IgD, CD27) **(C)** Frequency of RBD MBCs within the Spike MBCs population **(D)** Frequency of non-RBD MBCs within the Spike MBCs population **(E)** Affinity of RBD MBCs determined by the ratio MFI from the probe PE-CY7 divided by the MFI from immunoglobulin-associated beta (CD79b). **(F-H)** Frequency of RBD MBCs expressing IgG (F), IgM (G) or IgA (H) Means are plotted in the graphs. An unpaired, non-parametric Mann-Whitney test was used to compare the different groups. p-values are shown in the graphs and considered significant if lower than 0.05. **Sample size: treatment group n= 18 and placebo group n=11.**

### Association between MBC phenotype and viral load

Potential relationships between viral loads, mAb treatment, and MBC development were examined (**Fig. S2, S3A-H**). The frequency of CXCR3^+^ MBCs on day 28 in the placebo group correlated with viral RNA in both the nasal and nasopharyngeal swabs from post-infection (**Fig. S2B,D**). This correlation was lost for the mAb treatment group (**Fig. S2C,E**). These results suggest that CXCR3 expression on Spike MBCs is related to SARS-CoV-2 viral load at day 0 (study entry), for both nasal and nasopharyngeal viral loads.

### Impacts of later mRNA vaccination on SARS-CoV-2 MBCs

We then sought to understand whether, among SARS-CoV-2 infected individuals, previous anti-Spike mAb treatment would affect vaccination outcomes. A subset of the treatment cohort received two doses of immunization with original monovalent mRNA COVID-19 vaccines outside of the study with a median of 66 days post-treatment for the placebo group and 45 days post-treatment for the bamlanivimab group (**Fig. 3A**). At 24 weeks post-treatment, participants who received bamlanivimab and then were vaccinated had comparable frequencies of Spike MBCs (**Fig. 3B**), but significantly decreased frequencies of RBD MBCs (*p*=0.022, **Fig. 3C**) and they had decreased proportions of RBD MBCs among Spike MBCs (22.1% vs 10.3%, p<0.005, **Fig. 3D-E**) compared to placebo-treated vaccinated participants. The relative affinity of the RBD MBCs was also significantly lower in those previously treated with bamlanivimab and later vaccinated (*p*=0.034. **Fig. 3F**), suggesting that the treatment not only reduced the frequency but also the affinity maturation of RBD MBCs following vaccination. There were no RBD MBC isotype differences after vaccination in those treated with bamlanivimab compared to those who received placebo (**Fig. 3G-I**). In summary, the impact of mAb treatment on MBC frequencies and affinities was more significant after mRNA vaccine response compared to the endogenous response to infection.

**Figure 3.**
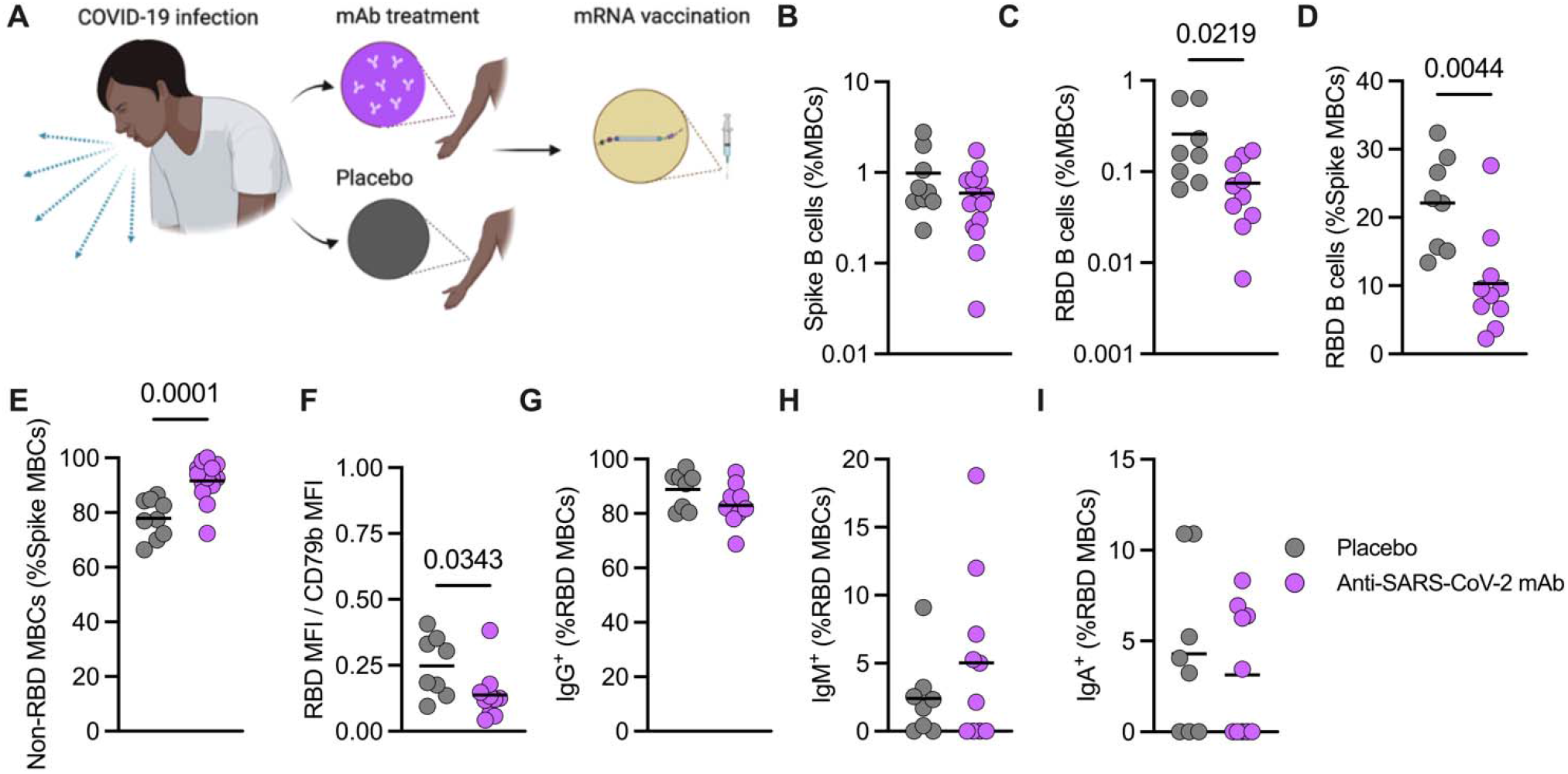
bamlanivimab treatment followed by mRNA vaccination decreases the frequency of RBD MBCs in the total MBC population and reduces affinity of RBD and Spike memory B cells. **(A)** Scheme of study design. COVID-19-infected humans receiving bamlanivimab were immunized with mRNA vaccines between 20 and 126 days after the mAb treatment. **(B-C)** Frequency of Spike (B) and RBD (C) MBCs among total memory B cell population **(D-E)** Frequency of RBD (D) and non-RBD (E) MBCs among the Spike MBCs population **(F)** Affinity of RBD MBCs determined by the ratio MFI from the RBDPE-CY7/ MFI from immunoglobulin-associated beta (CD79b). **(G-I)** Frequency of IgG (G), IgM (H) and IgA (I) RBD MBCs Means are plotted in the graphs. An unpaired, non-parametric Mann-Whitney test was used to compare the different groups. p-values are shown in the graphs and considered significant if lower than 0.05. **Sample size: treatment group n= 9, placebo group n=15**

### Memory B Cell Specificities

We assessed the impact of mAb treatment on MBCs in greater detail by examining RBD MBC epitope specificity. SARS-CoV-2 RBD and Spike antibody epitopes have been extensively characterized (22–25). We designed RBD epitope probe constructs by incorporating point mutations disrupting individual RBD epitopes, selected based on common SARS-CoV-2 escape mutations (**Fig. S6, Table 1**, **Methods**). Antibodies targeting the RBD have been divided into four major classes based on structural analyses of their epitopes (24, 26–32). Class I antibodies considerably overlap with the angiotensin-converting enzyme 2 (ACE2) receptor and bind when the RBD is in the open or up conformation (26–29). Class II antibodies have a partial overlap with the ACE2 receptor binding site. These are the most common antibodies elicited after COVID-19 infection (serum polyclonal antibodies) and mRNA vaccination. Bamlanivimab is classified as a class II site mAb.

**Table 1.** Mutations inserted in each of the 4 RBD constructs designed.

| Class of antibody binding | AA Changes |
| --- | --- |
| Class 1 | A417D, N460F, F486G |
| Class 2 | F456G, E484K, F490D |
| Class 3 | R346D, K444L, G446P |
| Class 4 | K417E, K378E |

Using RBD construct mutants corresponding to the antibody classes, we assessed the distribution of MBCs recognizing RBD sites I through IV, at 24 weeks post treatment. In bamlanivimab-treated unvaccinated individuals, the proportions of RBD MBCs targeting sites I through IV were not significantly different than the placebo group (**Fig. 4**). However, after vaccination in those previously treated with bamlanivimab, there was a significant shift away from MBCs recognizing the class II RBD epitope (*p* = 0.012, **Fig. 4**); the same epitope that is recognized by bamlanivimab. While the total frequency of Omicron BA.1 and BA.4.5 RBD MBCs was not significantly different between placebo and treatment groups, they were comparable to people who had received three vaccinations (uninfected donors, not enrolled in the ACTIV-2 trial) (**Fig. S4 D-F**). However, following treatment and vaccination, the proportion of cross-reactive WT RBD MBCs increased compared to placebo for BA.1 and BA.4.5 Omicron variants (**Fig. S4 G-I, Fig. S7**). Taken together, our data demonstrate that bamlanivimab treatment can alter B cell epitope recognition in response to SARS-CoV-2 mRNA vaccination.

**Figure 4.**
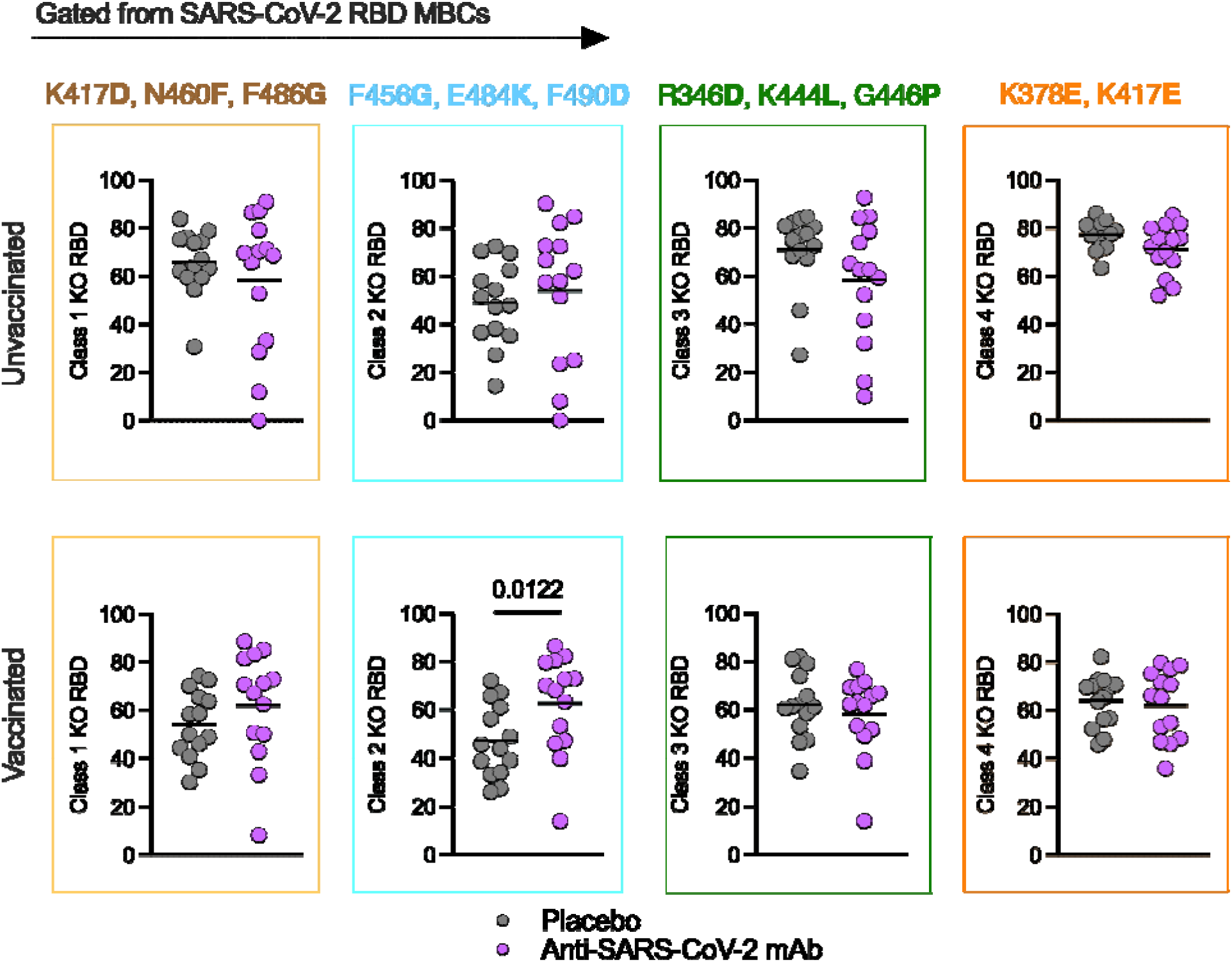
Bamlanivimab treatment followed by mRNA vaccination promotes escape of class II RBD MBCs. Graphs show the Frequency of B cells binding to RBD KO constructs 24 weeks post treatment. Means are plotted in th graphs. An unpaired, non-parametric Mann-Whitney test was used to compare the different groups. p-values are shown in th graphs and considered significant if lower than 0.05. **Sample size: vaccinated (treatment group n=14, placebo group n=14); unvaccinated (treatment group n=14, placebo group n=14).**

## DISCUSSION

The impact of mAb treatment in humans after antigen exposure, particularly during an ongoing immune response, has been unclear. Understanding how circulating antibodies from infection or vaccination, and also as a result of monoclonal antibody therapy, can affect SARS-CoV-2 B cell responses at the cellular level is valuable both for clinical considerations and for understanding B cell immunology. In this study, we evaluated Spike and RBD-specific MBCs from participants from the ACTIV-2 study who received a therapeutic mAb, bamlanivimab, during acute COVID-19 (4, 8, 20, 33). Our data demonstrate that bamlanivimab treatment altered epitope recognition by MBCs in SARS-CoV-2-infected people, indicating an antibody feedback phenomenon.

A previous study analyzed the memory B cell response in individuals who received two high-affinity anti-SARS-CoV-2 monoclonal antibodies and subsequently two doses of an mRNA vaccine (7). Specifically, they examinated MBC development in people without history of SARS-CoV-2 infection receiving two other anti-SARS-CoV-2 RBD monoclonal antibodies (C144-LS and C135-LS) and then vaccinated (7). The two study designs differ, but we observed multiple similarities in the immunological outcomes found between the two after mAb infusions: a) IgM+ MBCs were increased; b) RBD MBC affinities were reduced; and c) frequencies of MBCs specific for an RBD epitope recognized by the therapeutic mAb were reduced. Variant recognition can increase in response to vaccination (34, 35). Following vaccination, the frequency of MBCs that cross-reacted with BA.1 and BA.5 RBD was increased in the mAb treated group compared to the placebo group.

In a previous report, we had observed an association between SARS-CoV-2 infection and viral-vectored COVID-19 vaccination, and increased frequencies of CXCR3^+^ MBCs (36). Here, development of CXCR3^+^ MBCs were positively correlated with viral load in the respiratory mucosa a detected month prior B cell collection, indicating the CXCR3 expression by MBCs is likely programmed early during MBC development based on exposure to a cytokine mileu associated with viral infection. However, these data need to be interpreted considering that participants were enrolled in the study with different durations of symptoms, which could also impact on the viral load at entry.

Our work has limitations. The RBD constructs do not reflect the entirety of the SARS-CoV-2 escape spectrum. Additionally, the number of samples available to analyze the effect of bamlanivimab on the frequency of Omicron-binding MBCs was limited. Finally, mAbs were not isolated to directly measure Ab functionality of the SARS-CoV-2 RBD MBCs.

In sum, these findings are relevant for understanding antibody feedback phenomena, and for proper consideration of potential limitations of therapeutic mAb use, and challenges for universal flu and sarbecovirus development.

## Author Contributions

Conceptualization: CHC and SC. Investigation: CHC, NB, SIR; Formal Analysis: CHC and NB; Parent clinical trial design and data and sample collection: SFS, JR, CM, JJE, JSC, PK, DAW, ESD, JL, MDH, KWC, DMS; Binding antibody and virology assays: UMP, AH, SFS, AG; Data Curation: CHC; Writing: CHC and SC; Supervision: SC; Funding: SC, JSC.

## Competing Interest Statement

Disclose any competing interests here.

SC has consulted for GSK, Roche, Nutcracker Therapeutics, and Avalia. JSC has consulted for Merck and Company. PK is an employee and shareholder of Eli Lilly and Company. KWC has received research funding to the institution from Merck Sharp & Dohme and consulted for Pardes Biosciences. DMS has consulted for and has equity stake in Linear Therapies, Model Medicines, and Vx Biosciences and has consulted for Bayer, Kiadis, Signant Health, and Brio Clinical. The other authors do not declare any competing interests.

## Classification

Biological Sciences (Immunology)

## ACKNOWLEDGMENTS

The current work was funded by the National Institute of Allergy and Infectious Diseases (NIAID) of the National Institutes of Health (NIH) under award. This work was supported in part by the National Institute of Allergy and Infectious diseases (NIAID) of the National Institutes of Health (NIH), Department of Health and Human Services, under contract no. 75N93019C00065 (A.S, D.W.) and Collaborative Center from Human Immunology grant AI142742 (S.C.). Additional support was provided in part by the John and Mary Tu Foundation, NIH T32 AI007036 (S.I.R.), La Jolla Institute for Immunology institutional funds (S.C.), and an A.P. Giannini Foundation fellowship award (S.I.R.). Bamlanivimab was donated by Eli Lilly and Company. Lilly voluntarily asked the FDA to revoke the Emergency Use Authorization (EUA) for bamlanivimab 700 mg alone in April 2021. This request was not due to any new safety concerns. The content presented is the sole responsibility of the authors and may not represent the opinions of the NIH.

We would like to thank the study participants, site staff, site investigators, members of the ACTIV-2/A5401 Study Team, the ACTIV-2 Community Advisory Board, the AIDS Clinical Trials Group (including Lara Hosey, Jhoanna Roa, and Nilam Patel), the Harvard Center for Biostatistics in AIDS Research and ACTG Statistical and Data Analysis Center, the NIAID/Division of AIDS, Bill Erhardt, the ACTIV partnership (including Stacey Adams), and PPD for making this study possible.

## METHODS

### Study Population and Trial

ACTIV-2/A5401 is a multicenter phase 2/3 randomized controlled trial which was designed to evaluate the safety and efficacy of therapeutics for acute COVID-19 in non-hospitalized adults (4). Participants were enrolled in the United States between October-November 2020 (WT/ancestral SARS-CoV-2 infection period). Inclusion criteria included adults age 18 years or older with documented SARS-CoV-2 infection by FDA-authorized antigen- or molecular-based testing within seven days prior to study entry and no more than 10 days of symptoms at the time of enrollment. Participants were randomly assigned to the bamlanivimab (treatment) or placebo groups at a 1:1 ratio (referred to as the bamlanivimab cohort). Randomization was stratified by both time from COVID-19 symptom onset (less than or equal to 5 days post-symptom onset versus greater than 5 days) and risk of progression to severe COVID-19 (low versus high, based on age and comorbid medical conditions) (4). Additional information regarding the ACTIV-2/A5401 clinical trial is available at ClinicalTrials.gov (Identifier: NCT04518410) and can be found in the supplementary materials for the primary outcomes manuscript (4). IgG titers for a subset of study participants have been previously reported (8).

PBMC samples from selected participants who received bamlanivimab 700 mg or its placebo were available for this study (**S1 Table**). Participants in this study were representative of the larger ACTIV-2/A5401 bamlanivimab cohort, including risk for progression to severe COVID-19, baseline serostatus, and time from COVID-19 symptom onset to enrollment (8). The number of samples tested in each experiment is described in the figure legends.

PBMCs from uninfected and 3-times vaccinated donors were obtained with IRB approval from the La Jolla Institute for Immunology Healthy Donor Program.

### Nasal and nasopharyngeal swab collection and quantification of SARS-CoV-2 RNA

Anterior nasal and nasopharyngeal (NP) swab samples were collected and processed using standardized reagents and procedures, as previously reported (4). Nasal swabs were collected on site during in person study visits at day 28. Nasal and NP swab samples collected on site were frozen at the same day as collection. Nasal swab samples collected remotely were kept chilled until returned to the study site, and then frozen upon arrival at the study site. All nasal swabs were frozen within 7 days of collectionStudy day 0 nasal and NP swab samples were collected prior to treatment. RNA extraction, reverse transcription, amplification, and quantification of SARS-CoV-2 RNA was performed at a centralized laboratory (University of Washington) using an FDA-approved quantitative reverse transcription polymerase chain reaction assay. The limit of detection (LOD) for SARS-CoV-2 RNA for this assay was 1.4 log10 copies/mL.. For samples with viral RNA levels above the ULOQ, samples were diluted, and the assay was repeated to obtain a quantitative value.

### Serology

Serum binding antibody assays were performed to evaluate IgG responses to SARS-CoV-2 nucleocapsid and ancestral spike S2 domains and RBD using the Bio-Plex Pro Human SARS-CoV-2 IgG (N, S2, RBD) 4-Plex Panel serology assay (Bio-Rad 12014634) as previously described (8).

### RBD constructs design

We designed four SARS-CoV-2 RBD biotinylated constructs containing the most relevant mutations leading to an escape of monoclonal antibodies from each of the classes I to IV (**Table 1**), according to the scientific literature available (26, 28). These constructs were expressed by Acrobio Inc. Their identity was confirmed by ELISA using a panel of mAbs.

### SARS-CoV-2 B cell probe staining and flow cytometry assay

Before use, cryopreserved PBMC were thawed at 37°C and then resuspended in pre-warmed complete RPMI medium with 10% human AB serum (Gemini Bioproducts) benzonase. Cell counts and viability were assessed on a Muse Cell Analyzer (Luminex) using the Muse Count & Viability Kit after washing. SARS-CoV-2-specific memory B cells were detected using B cell probes as previously described (13, 37). Briefly, biotinylated full-length SARS-CoV-2 Spike protein was purchased from Acro Biosystems and SARS-CoV-2 Spike protein Receptor-Binding Domain (RBD) from BioLegend. SARS-CoV-2 Spike protein Receptor-Binding Domain (RBD) variants were also purchased from Acro Biosystems. To enhance specificity, both ancestral Spike- and RBD-specific MBCs were identified using two fluorochromes for each protein. Thus, the biotinylated SARS-CoV-2 spike was multimerized with either Streptavidin Alexa Fluor 647 or BV421 at a 20:1 ratio (6:1 M ratio) for 1 hour 4°C. Biotinylated RBD was multimerized with Streptavidin BV711 or PE-Cy7 at a 2.2:1 ratio (4:1 M ratio). Streptavidin PE-Cy5.5 was used as a decoy probe to remove any SARS-CoV-2 nonspecific streptavidin-binding B cells. Then, PBMCs were placed in U-bottom 96 well plates and stained with a solution consisting of 5mM of biotin to avoid cross-reactivity among probes, 20 ng of decoy probe, 150 ng of Spike per probe and 16.4 ng of RBD per probe per sample (300 ng total Spike and 32.8 ng total RBD per sample), diluted in Brilliant Buffer and incubated for 1 hour at 4°C, protected from light. After washing with FACS buffer (PBS with 2% FBS, NaN3, EDTA), cells were incubated with surface antibodies diluted in Brilliant Staining Buffer for 30 min at 4C in the dark. Viability staining was performed using Live/Dead Fixable Blue Stain Kit diluted at 1:200 in DPBS (Mg^2+^/Ca^2+^) and incubated at 4 °C for 30 min. Cells were washed twice in DPBS and resuspended in FACS buffer before acquisition. The acquisition was performed using Cytek Aurora. The frequency of antigen-specific MBCs was expressed as a percentage of total memory B cells (Live, Lymphocytes, Singlets, CD3^−^ CD14^−^ CD16^−^ CD56^−^CD19^+^, CD19^+^ CD20^+^, CD20^+^ CD38^int/–^, IgD^−^ and/or CD27^+^).

### Statistical analyses

Statistical analyses were performed using GraphPad version 9. Comparisons between groups were performed using Mann-Whitney tests. P values are shown in the graphs and are considered significant if lower than 0.05.

## REFERENCES

1. RECOVERY Collaborative Group, Casirivimab and imdevimab in patients admitted to hospital with COVID-19 (RECOVERY): a randomised, controlled, open-label, platform trial. Lancet Lond. Engl. 399, 665–676 (2022).

2. D. M. Weinreich, et al., REGN-COV2, a Neutralizing Antibody Cocktail, in Outpatients with Covid-19. N. Engl. J. Med. 384, 238–251 (2021).

3. M. Dougan, et al., Bamlanivimab plus Etesevimab in Mild or Moderate Covid-19. N. Engl. J. Med. 385, 1382– 1392 (2021).

4. K. W. Chew, et al., Antiviral and clinical activity of bamlanivimab in a randomized trial of non-hospitalized adults with COVID-19. Nat. Commun. 13, 4931 (2022).

5. J. Boucau, et al., Monoclonal antibody treatment drives rapid culture conversion in SARS-CoV-2 infection. Cell Rep. Med. 3, 100678 (2022).

6. R. J. Benschop, et al., The anti-SARS-CoV-2 monoclonal antibody bamlanivimab minimally affects the endogenous immune response to COVID-19 vaccination. Sci. Transl. Med. 14, eabn3041 (2022).

7. D. Schaefer-Babajew, et al., Antibody feedback regulates immune memory after SARS-CoV-2 mRNA vaccination. Nature 613, 735–742 (2023).

8. S. I. Ramirez, et al., Bamlanivimab therapy for acute COVID-19 does not blunt SARS-CoV-2-specific memory T cell responses. JCI Insight 7, e163471 (2022).

9. B. Heyman, Regulation of antibody responses via antibodies, complement, and Fc receptors. Annu. Rev. Immunol. 18, 709–737 (2000).

10. J. M. J. Tas, et al., Antibodies from primary humoral responses modulate the recruitment of naive B cells during secondary responses. Immunity 55, 1856–1871.e6 (2022).

11. K. A. Pape, J. J. Taylor, R. W. Maul, P. J. Gearhart, M. K. Jenkins, Different B cell populations mediate early and late memory during an endogenous immune response. Science 331, 1203–1207 (2011).

12. T. Smith, ACTIVE IMMUNITY PRODUCED BY SO CALLED BALANCED OR NEUTRAL MIXTURES OF DIPHTHERIA TOXIN AND ANTITOXIN. J. Exp. Med. 11, 241–256 (1909).

13. Y. Zhang, et al., Germinal center B cells govern their own fate via antibody feedback. J. Exp. Med. 210, 457– 464 (2013).

14. W. B. Alsoussi, et al., SARS-CoV-2 Omicron boosting induces de novo B cell response in humans. Nature 617, 592–598 (2023).

15. C. I. Kaku, et al., Evolution of antibody immunity following Omicron BA.1 breakthrough infection. Nat. Commun. 14, 2751 (2023).

16. C. I. Kaku, et al., Recall of preexisting cross-reactive B cell memory after Omicron BA.1 breakthrough infection. Sci. Immunol. 7, eabq3511 (2022).

17. M. Koutsakos, A. H. Ellebedy, Immunological imprinting: Understanding COVID-19. Immunity 56, 909–913 (2023).

18. H. L. Dugan, et al., Preexisting immunity shapes distinct antibody landscapes after influenza virus infection and vaccination in humans. Sci. Transl. Med. 12, eabd3601 (2020).

19. A. Schiepers, et al., Molecular fate-mapping of serum antibody responses to repeat immunization. Nature 615, 482–489 (2023).

20. M. C. Choudhary, et al., Emergence of SARS-CoV-2 escape mutations during Bamlanivimab therapy in a phase II randomized clinical trial. Nat. Microbiol. 7, 1906–1917 (2022).

21. K. A. Pape, et al., High-affinity memory B cells induced by SARS-CoV-2 infection produce more plasmablasts and atypical memory B cells than those primed by mRNA vaccines. Cell Rep. 37, 109823 (2021).

22. P. J. M. Brouwer, et al., Potent neutralizing antibodies from COVID-19 patients define multiple targets of vulnerability. Science 369, 643–650 (2020).

23. K. M. Hastie, et al., Defining variant-resistant epitopes targeted by SARS-CoV-2 antibodies: A global consortium study. Science 374, 472–478 (2021).

24. C. Fan, et al., Neutralizing monoclonal antibodies elicited by mosaic RBD nanoparticles bind conserved sarbecovirus epitopes. Immunity 55, 2419–2435.e10 (2022).

25. D. F. Robbiani, et al., Convergent antibody responses to SARS-CoV-2 in convalescent individuals. Nature 584, 437–442 (2020).

26. A. J. Greaney, et al., Mapping mutations to the SARS-CoV-2 RBD that escape binding by different classes of antibodies. Nat. Commun. 12, 4196 (2021).

27. A. J. Greaney, et al., Comprehensive mapping of mutations in the SARS-CoV-2 receptor-binding domain that affect recognition by polyclonal human plasma antibodies. Cell Host Microbe 29, 463–476.e6 (2021).

28. A. J. Greaney, et al., Complete Mapping of Mutations to the SARS-CoV-2 Spike Receptor-Binding Domain that Escape Antibody Recognition. Cell Host Microbe 29, 44–57.e9 (2021).

29. A. J. Greaney, T. N. Starr, J. D. Bloom, An antibody-escape estimator for mutations to the SARS-CoV-2 receptor-binding domain. Virus Evol. 8, veac021 (2022).

30. Z. Wang, et al., mRNA vaccine-elicited antibodies to SARS-CoV-2 and circulating variants. Nature 592, 616– 622 (2021).

31. A. Addetia, et al., Structural changes in the SARS-CoV-2 spike E406W mutant escaping a clinical monoclonal antibody cocktail. Cell Rep. 42, 112621 (2023).

32. A. R. Shiakolas, et al., Efficient discovery of SARS-CoV-2-neutralizing antibodies via B cell receptor sequencing and ligand blocking. Nat. Biotechnol. 40, 1270–1275 (2022).

33. T. N. Starr, A. J. Greaney, A. S. Dingens, J. D. Bloom, Complete map of SARS-CoV-2 RBD mutations that escape the monoclonal antibody LY-CoV555 and its cocktail with LY-CoV016. Cell Rep. Med. 2, 100255 (2021).

34. K. Röltgen, et al., Immune imprinting, breadth of variant recognition, and germinal center response in human SARS-CoV-2 infection and vaccination. Cell 185, 1025–1040.e14 (2022).

35. P. Sun, et al., Asymptomatic or symptomatic SARS-CoV-2 infection plus vaccination confers increased adaptive immunity to variants of concern. iScience 25, 105202 (2022).

36. Z. Zhang, et al., Humoral and cellular immune memory to four COVID-19 vaccines. Cell 185, 2434–2451.e17 (2022).

37. A. Tarke, et al., SARS-CoV-2 vaccination induces immunological T cell memory able to cross-recognize variants from Alpha to Omicron. Cell 185, 847–859.e11 (2022).

