## Supplementary material for "SARS-CoV-2 monoclonal antibody treatment followed by vaccination shifts human memory B cell epitope recognition suggesting antibody feedback"

**Figures**

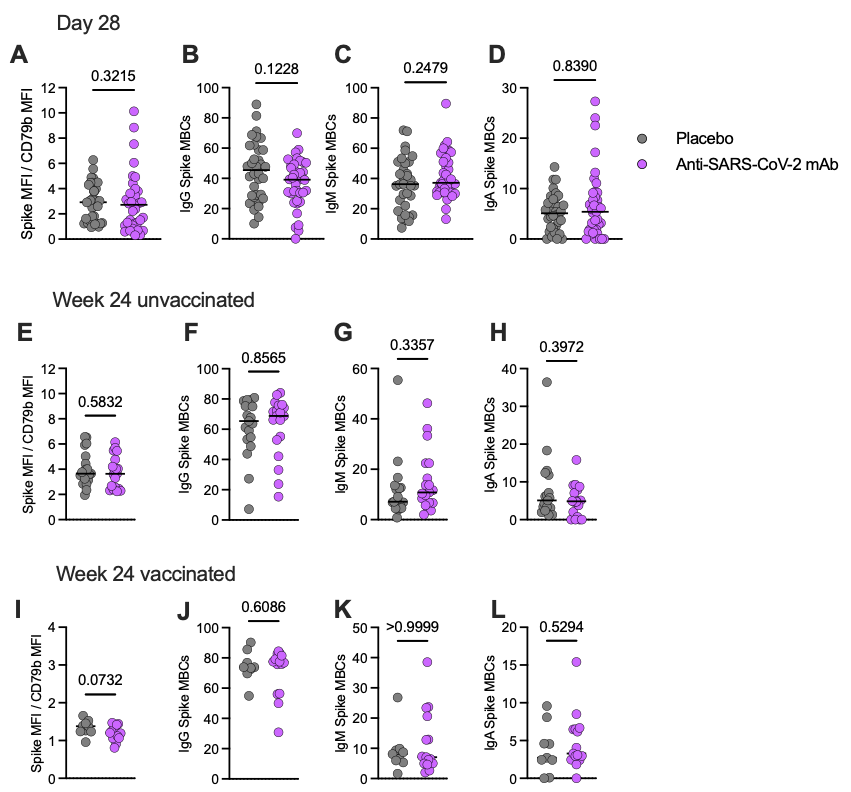

**S1 Fig. Spike MBCs isotypes and Spike MBCs affinity in the timepoints analyzed.**

Affinity was measured by the ratio of Spike MFI/CD79b. IgG, IgM and IgA frequencies were assessed.

**(A-D)** Day 28 post mAb treatment (n=25 placebo and n=21 treatment).

**(E-H)** Week 24 post mAb treatment in unvaccinated donors (n=18 placebo and n=11 treatment).

**(I-L)** Week 24 post mAb treatment and further mRNA vaccinated donors (n=9 placebo and n=15 treatment).

Comparisons were performed using the Mann-Whitney test (**p* < 0.05, ***p* <0.01, ****p* < 0.001).

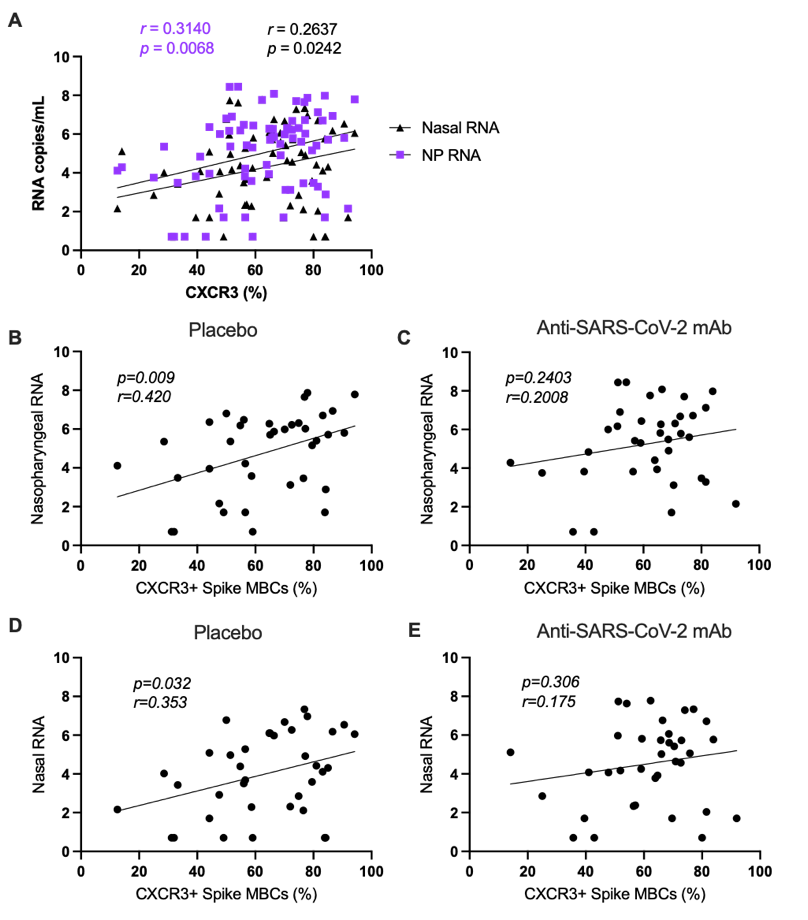

**Figure S2. Correlations between nasal and nasopharyngeal viral load (day 0/ study entry) and frequency of CXCR3-expressing MBCs (day 28)**

**(A).** Pearson correlation between viral RNA load in nasal tissue and nasopharyngeal tissue with all donors (treatment n=21 and placebo n=25) included

**(B-C).** Pearson correlation between viral RNA load in the nasopharyngeal tissue of (B) placebo and (C) mAb-treated donors

**(D-E).** Pearson correlation between viral RNA load in the nasal tissue of (A) placebo and (B) mAb-treated donors

*p*-values: **p* < 0.05, ***p* <0.01, ****p* < 0.001.

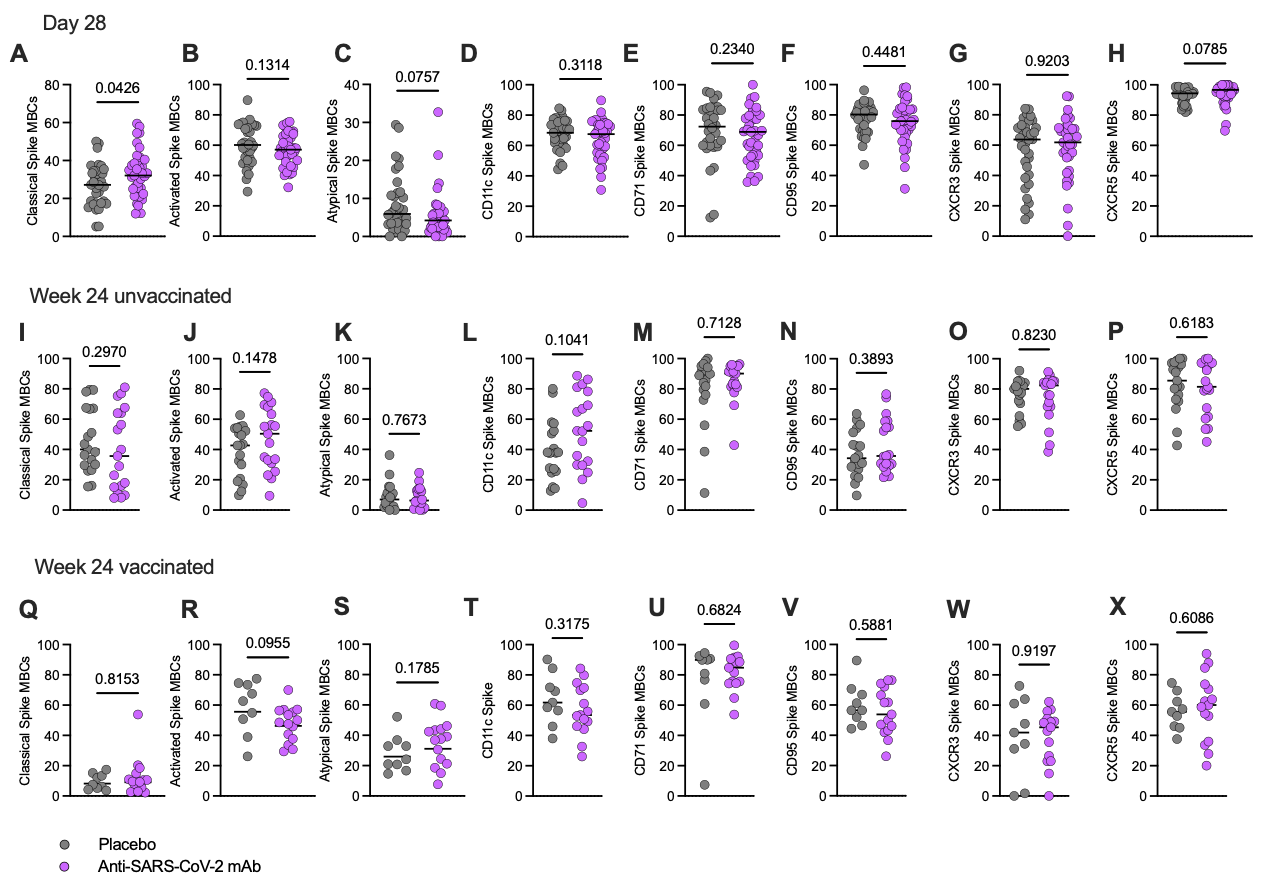

**Figure S3. Phenotypic characterization of B cells after mAb treatment**.

Classical, activated, and atypical memory B cells and the frequency of memory B cells expressing CD11c, CD71, CD95, CXCR3, and CXCR5 are shown.

**(A-H)** Day 28 post mAb treatment (n=25 placebo and n=21 treatment).

**(I-P)** Week 24 post mAb treatment in unvaccinated donors (n=25 placebo and n=21 treatment).

**(Q-X)** Week 24 post mAb treatment and further mRNA vaccinated donors (n= 9 placebo and n=15 treatment).

Comparisons were performed using the Mann-Whitney test (**p* < 0.05, ***p* <0.01, ****p* < 0.001).

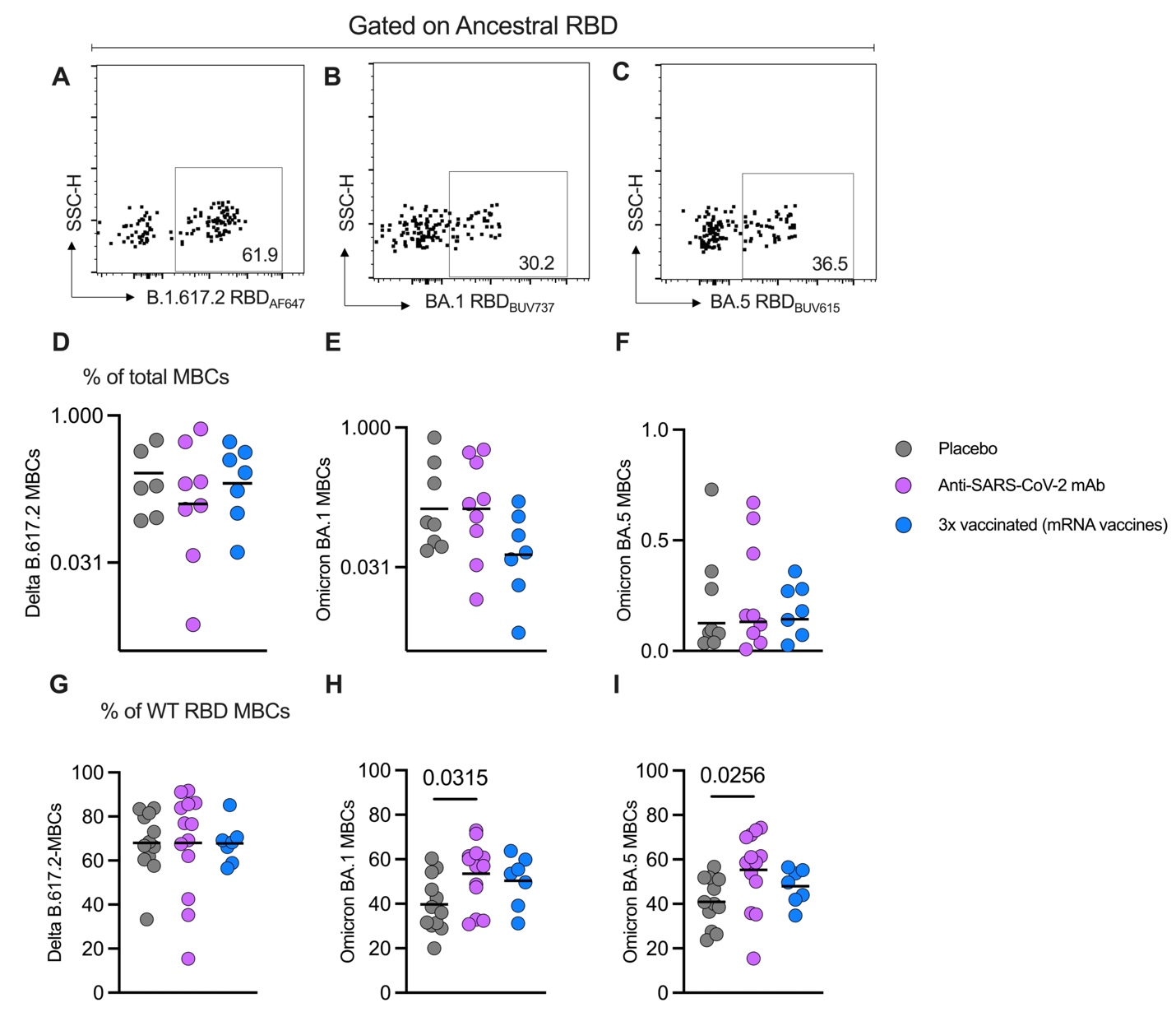

**Figure S4. mRNA vaccination followed by Bamlanivimab treatment increases the frequency of Omicron BA.1 and BA.5- specific MBCs to a level comparable to 3x mRNA-vaccinated donors**

**A-C.** Representative flow cytometry gating strategy to select Delta, Omicron BA.1 and BA.5 RBD-specific MBCs

**D-F.** Frequency of Delta, Omicron BA.1 and BA.5 RBD-specific MBCs in the placebo, treatment and in comparison, with donors receiving 3 doses of mRNA vaccines.

**G-F.** Same cells as **D-F**, but gated as % of Spike MBCs.

Kruskal-Wallis test was used as a statistical test to compare different groups.

Comparisons were performed using the Kruskal-Wallis test followed by Dunn’s multiple comparisons test (**p* < 0.05, ***p* <0.01, ****p* < 0.001).

Sample size: placebo n=12, treatment n=13, 3x vaccinated=7

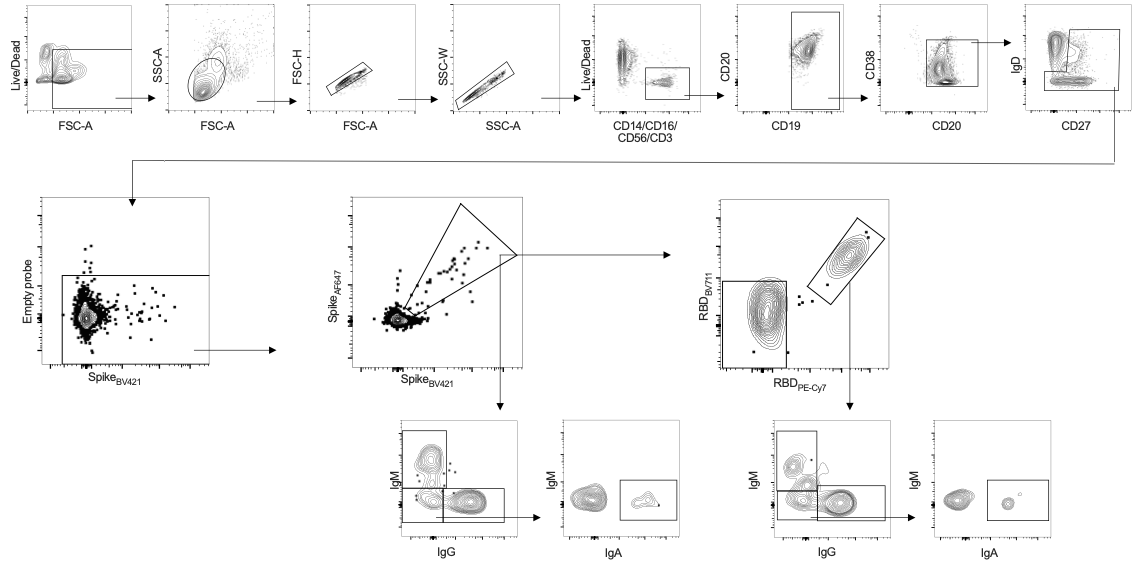

**Figure S5. Flow cytometry gating strategy to identify and quantify SARS-CoV-2 (Spike and RBD) -specific memory B cells.**

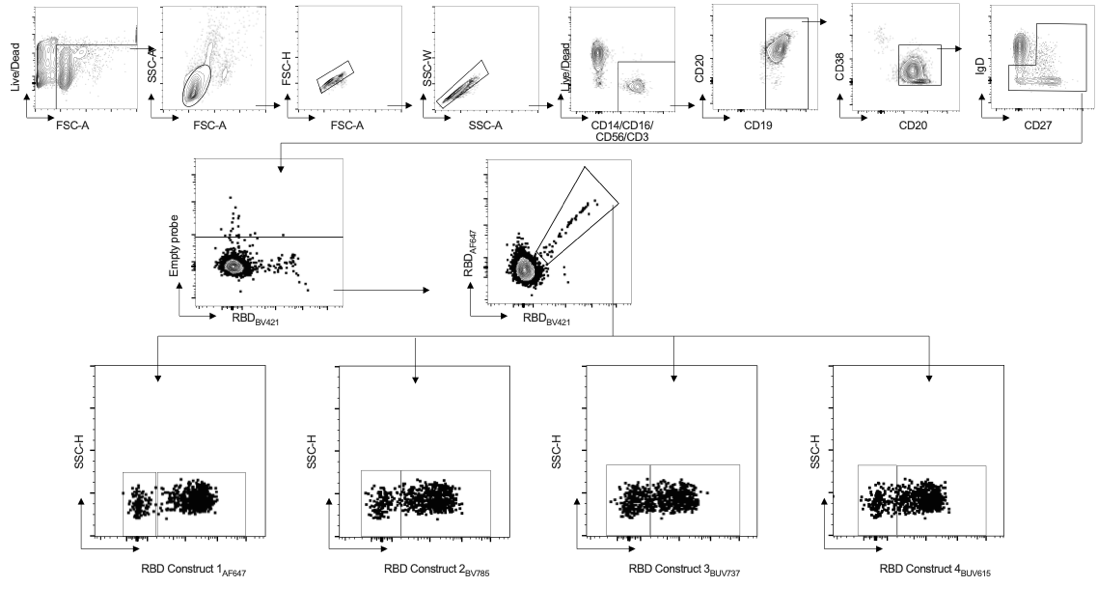

**Figure S6. Flow cytometry gating strategy to identify and quantify memory B cells binding the RBD constructs containing the most frequent mutations escaping monoclonal antibodies from classes I to IV.**

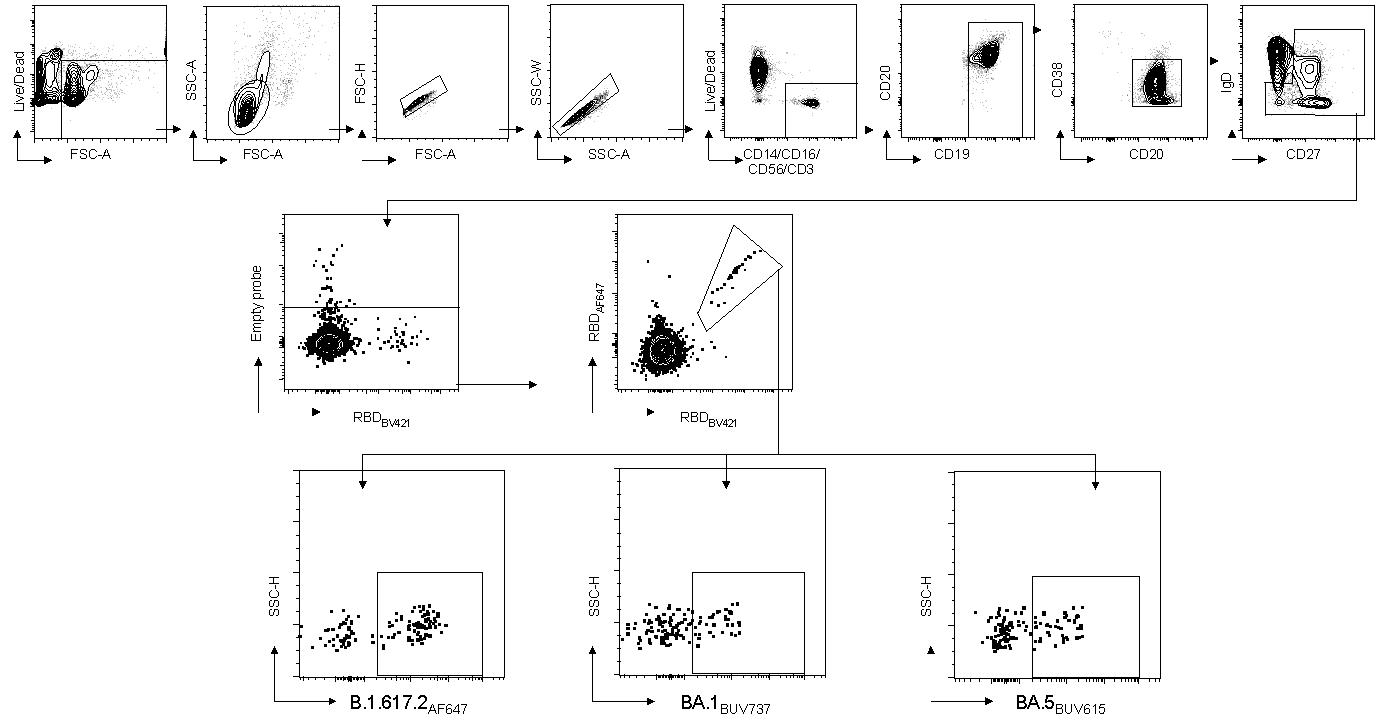

**Figure S7. Flow cytometry gating strategy to identify and quantify memory B cells binding to SARS-CoV-2 variants of concern (Delta, Omicron BA.1 and Omicron BA.5).**

**Tables**

**S1 Table. mAb Treatment and Placebo group participant characteristics**

|  | **Bamlanivimab (n = 46)** | **Placebo (n = 49)** |
| --- | --- | --- |
| **Age (years)** | 18-86 [Median = 46, IQR =22] | 19-72 [Median =43, IQR = 20.5] |
| **Sex** |  |  |
| Male | 59% (27/46) | 49% (24/49) |
| Female | 40% (19/46) | 51% (25/49) |
| **Race** |  |  |
| African American  or Black | 15% (7/46) | 10% (5/49) |
| Alaskan Native or  American Indian | 2% (1/46) | 0% (0/49) |
| Asian | 2% (1/46) | 6% (3/49) |
| Native Hawaiian or  Pacific Islander | 0% (0/46) | 0% (0/49) |
| Other/Mixed Race | 2% (1/46) | 6% (3/49) |
| Unknown | 0% (0/46) | 2% (1/49) |
| White | 78% (36/46) | 76% (37/49) |
| **Ethnicity** |  |  |
| Hispanic | 13% (6/46) | 24% (12/49) |
| Non-Hispanic | 87% (42/46) | 65% (35/49) |
| Unknown | 0% (0/46) | 4% (2/49) |
| **Sample Collection Dates** | October-November 2020 | October-November 2020 |
| **SARS-CoV-2 PCR Positivity** | 100% (46/46) | 100% (49/49) |
| **Days post-symptom onset at randomization** |  |  |
| ≤ 5 days | 30% (14/46) | 27% (13/49) |
| > 5 days | 70% (32/46) | 73% (36/49) |
| **Baseline serostatus** |  |  |
| Seropositive (by RBD IgG) | 30% (14/46) | 29% (14/49) |
| Seronegative (by RBD IgG) | 70% (32/46) | 71% (35/49) |
| **Vaccination status** |  |  |
| Vaccinated at entry | 0% (0/46) | 0% (0/49) |
| Vaccinated at day 28 | 0% (0/46) | 0% (0/49) |
| **Risk group for severe COVID-19** |  |  |
| High risk group | 52% (24/46) | 49% (24/49) |
| Low risk group | 48% (22/46) | 51% (25/49) |

**S2 Table. Antibodies used for flow cytometry staining of MBCs.**

| **Reagent** | **Source** | **Identifier** | **Dilution** |
| --- | --- | --- | --- |
| Live/Dead Fixable Blue Stain Kit | Thermofisher | #L34962 | 1:200 |
| Mouse anti-human CD19 BUV563 (SJ25C1) | BD Bioscience | #612916 | 1:200 |
| Mouse anti-human CD95 BUV737 (clone DX2) | BD Bioscience | #612790 | 1:200 |
| Mouse anti-human CD183 (CXCR3) BV605 (clone G025H7) | Biolegend | #353728 | 1:50 |
| Mouse anti-human IgD Pacific Blue (clone IA6-2) | Biolegend | #348224 | 1:50 |
| Mouse anti-human CD20 Brilliant Violet 510 (clone 2H7) | Biolegend | #302340 | 1:100 |
| Mouse anti-human IgM Brilliant Violet 570 (clone MHM-88) | Biolegend | #314517 | 1:200 |
| Mouse anti-human CD27 BB515 (clone M-T271) | BD Bioscience | #564642 | 1:200 |
| Mouse anti-human IgA Vio Bright FITC (clone IS11-8E10) | Miltenyi Biotec | #130-113-480 | 1:400 |
| Mouse anti-human CD3 PerCP (clone SK7) | Biolegend | #344814 | 1:100 |
| Mouse anti-human CD14 PerCP (clone 63D3) | Biolegend | #367152 | 1:200 |
| Mouse anti-human CD16 PerCP (clone 3G8) | Biolegend | #302030 | 1:200 |
| Mouse anti-human CD56 PerCP (clone 3G8) | Biolegend | #318342 | 1:200 |
| Mouse anti-human IgG PerCP/Cyanine5.5 (clone M1310G05) | Biolegend | #410710 | 1:100 |
| Mouse anti-human CD71 PE/Dazzle 594 (clone CY1G4) | Biolegend | #334120 | 1:200 |
| Mouse anti-human CD11c PE/Cy5 (clone 3.9) | Biolegend | #301610 | 1:200 |
| Mouse anti-human CD21 Alexa Fluor 700 (clone Bu32) | Biolegend | #354918 | 1:50 |
| Mouse anti-human CD38 APC/Fire 810 (clone HIT2) | Biolegend | #303550 | 1:200 |
| Mouse anti-human CD79b PE (clone CB3-1) | Biolegend | #341404 | 1:200 |
| Rat anti-human CD185 (CXCR5) BV750 (clone RF8B2) | BD Bioscience | #747111 | 1:200 |
| Brilliant violet 421 Streptavidin | Biolegend | #405225 |  |
| Brilliant violet 711 Streptavidin | Biolegend | #405241 |  |
| Brilliant violet 785 Streptavidin | Biolegend | #405249 |  |
| Streptavidin, Alexa Fluor 647 conjugate | invitrogen | #S21374 |  |
| BD Horizon BUV615 Streptavidin | BD Biosciences | #613013 |  |
| BD Horizon BUV737 Streptavidin | BD Biosciences | #612775 |  |
| Streptavidin PE-Cy 5.5 Conjugate | invitrogen | #SA1018 |  |

**Supplemental Acknowledgments**

**ACTIV-2/A5401 Study Team**

- Kara Chew, MD, MS, Co-Chair, David Geffen School of Medicine at University of California, Los Angeles, Los Angeles, CA, USA
- David (Davey) Smith, MD, MAS, Co-Chair, University of California, San Diego, La Jolla, CA, USA
- Eric Daar, MD, Vice Chair, Lundquist Institute at Harbor-UCLA Medical Center, Torrance, CA, USA
- David Wohl, MD, Vice Chair, University of North Carolina at Chapel Hill School of Medicine, Chapel Hill NC, USA
- Judith Currier, MD, MSc, Protocol Investigator and ACTG Chair, David Geffen School of Medicine at University of California, Los Angeles, Los Angeles, CA, USA
- Joseph Eron, MD, Protocol Investigator and ACTG Vice Chair, University of North Carolina at Chapel Hill School of Medicine, Chapel Hill NC, USA
- Arzhang Cyrus Javan, MD, MPH, DTM&H, NIH Division of AIDS (DAIDS) Clinical Representative, National Institutes of Health, Rockville, MD, USA
- Michael Hughes, PhD, Lead Statistician, Harvard T.H. Chan School of Public Health, Boston, MA, USA
- Carlee Moser, PhD, Statistician, Harvard T.H. Chan School of Public Health, Boston, MA, USA
- Mark Giganti, PhD, Statistician, Harvard T.H. Chan School of Public Health, Boston, MA, USA
- Justin Ritz, MS, Statistician, Harvard T.H. Chan School of Public Health, Boston, MA, USA
- Lara Hosey, MA, Clinical Trials Specialist, AIDS Clinical Trials Group (ACTG) Network Coordinating Center, Social & Scientific Systems, a DLH Company, Silver Spring, MD, USA
- Jhoanna Roa, MD, Clinical Trials Specialist, AIDS Clinical Trials Group (ACTG) Network Coordinating Center, Social & Scientific Systems, a DLH Company, Silver Spring, MD, USA
- Nilam Patel, Clinical Trials Specialist, AIDS Clinical Trials Group (ACTG) Network Coordinating Center, Social & Scientific Systems, a DLH Company, Silver Spring, MD, USA
- Kelly Colsh, PharmD, DAIDS Pharmacist, NIH/DAIDS Pharmaceutical Affairs Branch, Rockville, MD, USA
- Irene Rwakazina, PharmD, DAIDS Pharmacist, NIH/DAIDS Pharmaceutical Affairs Branch, Rockville, MD, USA
- Justine Beck, PharmD, DAIDS Pharmacist, NIH/DAIDS Pharmaceutical Affairs Branch, Rockville, MD, USA
- Scott Sieg, PhD, Protocol Immunologist, Case Western Reserve University, Cleveland, OH, USA
- Jonathan Li, MD, MMSc, Protocol Virologist, Brigham and Women’s Hospital, Harvard Medical School, Boston, MA, USA
- Courtney Fletcher, PharmD, Protocol Pharmacologist, University of Nebraska Medical Center, Omaha, NE, USA
- William Fischer MD, Protocol Critical Care Specialist, University of North Carolina at Chapel Hill School of Medicine, Chapel Hill NC, USA
- Teresa Evering, MD, MS, Protocol Investigator, Weill Cornell Medicine, New York, NY, USA
- Rachel Bender Ignacio, MD, MPH, Protocol Investigator, University of Washington, Seattle, WA, USA
- Sandra Cardoso, MD, PhD, Protocol Investigator, Fundação Oswaldo Cruz, Rio de Janeiro, Brazil
- Katya Corado, MD, Lundquist Institute at Harbor-UCLA Medical Center, Torrance, CA, USA
- Prasanna Jagannathan, MD, Protocol Investigator, Stanford University School of Medicine, Palo Alto, CA, USA
- Nikolaus Jilg, MD, PhD, Protocol Investigator, Massachusetts General Hospital, Harvard Medical School, Boston, MA, USA
- Alan Perelson, PhD, Protocol Investigator, Los Alamos National Laboratory, Los Alamos, NM, USA
- Sandy Pillay, MB, CHB, Protocol Investigator, Enhancing Care Foundation, Durban, KwaZulu- Natal, South Africa
- Cynthia Riviere, MD, Protocol Investigator, GHESKIO Center, Port-au-Prince, Haiti
- Upinder Singh, MD, Protocol Investigator, Stanford University School of Medicine, Palo Alto, CA, USA
- Babafemi Taiwo, MBBS, MD, Protocol Investigator, Northwestern University Feinberg School of Medicine, Chicago, IL, USA
- Joan Gottesman, BSN, RN, CCRP, Field Representative, Vanderbilt University Medical Center, Nashville, TN, USA
- Matthew Newell, BSN, RN, CCRN, Field Representative, University of North Carolina at Chapel Hill School of Medicine, Chapel Hill NC, USA
- Susan Pedersen, BSN, RN, Field Representative, University of North Carolina at Chapel Hill School of Medicine, Chapel Hill NC, USA
- Joan Dragavon, MLM, Laboratory Technologist, University of Washington, Seattle, WA, USA
- Cheryl Jennings, BS, Laboratory Technologist, Northwestern University, Chicago, IL, USA Brian Greenfelder, BA, Laboratory Technologist, Ohio State University, Columbus, OH, USA
- William Murtaugh, MPH, Laboratory Specialist, ACTG Laboratory Center, University of California, Los Angeles, Los Angeles, CA, USA
- Jan Kosmyna, MIS, RN, CCPR, ACTG Community Scientific Subcommittee Representative, Case Western University Clinical Research Site, North Royalton, OH, USA
- Morgan Gapara, MPH, International Site Specialist, ACTG Network Coordinating Center, Social & Scientific Systems, a DLH Company, Durham, NC, USA
- Akbar Shahkolahi, PhD, International Site Specialist, ACTG Network Coordinating Center, Social & Scientific Systems, a DLH Company, Silver Spring, MD, USA
- Paul Klekotka, MD, PhD, Industry Representative, Lilly Research Laboratories, San Diego, CA, USA
